# Deep learning model for ultrafast quantification of blood flow in diffuse correlation spectroscopy

**DOI:** 10.1101/2020.06.24.167882

**Authors:** Chien-Sing Poon, Feixiao Long, Ulas Sunar

**Author notes:** Corresponding author: Ulas Sunar.

## Abstract

Diffuse correlation spectroscopy (DCS) is increasingly used in the optical imaging field to assess blood flow in humans due to its non-invasive, real-time characteristics and its ability to provide label-free, bedside monitoring of blood flow changes. Previous DCS studies have utilized a traditional curve fitting of the analytical or Monte Carlo models to extract the blood flow changes, which are computationally demanding and less accurate when the signal to noise ratio decreases. Here, we present a deep learning model that eliminates this bottleneck by solving the inverse problem more than 2300% faster, with equivalent or improved accuracy compared to the nonlinear fitting with an analytical method. The proposed deep learning inverse model will enable real-time and accurate tissue blood flow quantification with the DCS technique.

## INTRODUCTION

Blood flow has been investigated as an important biomarker for a variety of diseases in humans. Most diseases induce changes in blood flow thus blood flow parameter can be indicative of the diseases. Blood flow has also been an indicator of muscle activation, where the muscle fibers increase the rate of oxygen consumption and therefore increase the rate of oxygen supply by increasing blood flow [1,2]. Several clinical technologies for assessing tissue blood flow are available, including computed tomography and magnetic resonance imaging. However, these modalities provide single snapshot time points and are not suited for continuous, long-term monitoring at the bedside settings [3]. Laser Doppler flowmetry measures blood flow from a very small volume, while Doppler ultrasound measures blood flow in a large vasculature, and it is hard to use for longitudinal monitoring due to the need for stable ultrasonic probe orientation [3].

Near-infrared diffuse correlation spectroscopy (DCS), also known as diffusing-wave spectroscopy (DWS), is an emerging technique for measuring blood flow in humans. It uses the well-known spectral window in the near-infrared (NIR, 650 to 900 nm) range, wherein the relatively low biological tissue absorption enables deep penetration of light. Being a label-free technique, DCS is highly suitable for continuous noninvasive measurement of blood flow in biological tissues. In the last decade, DCS technology has been implemented, extensively validated, and employed to measure blood flow noninvasively in deep tissues, such as muscle and brain functional studies [4–9]. Recently, by using DCS we have shown the evidence of assessing resting-state functional connectivity between brain regions in humans by inspecting low-level oscillations in the blood flow changes [10,11].

The current quantification approach of DCS is to fit iteratively the measured intensity autocorrelation function with either the analytical or the Monte Carlo model. This step can be time-consuming, especially for dynamic longitudinal measurements, thus it is typically performed as a post-processing step. However, it is useful to obtain the results in real-time to provide feedback to the clinicians. The deep learning method allows fast retrieval of parameters and been used recently in the diffuse optical field, including functional near-infrared spectroscopy [12–14], spatial frequency domain imaging [15], fluorescence imaging and tomography [16–18]. Its application for DCS has yet to be established [19]. In this paper, we show the deep learning implementation of DCS to quantify blood flow-related parameter and we apply this implementation for the assessment of blood flow changes during arm cuff ischemia protocol on a human subject.

## MATERIALS AND METHODS

### DIFFUSE CORRELATION SPECTROSCOPY

In this short communication, we are not going into the details of the diffuse correlation spectroscopy (DCS) technique, which can be found in recent reviews [6,9,20,21]. Briefly, the normalized diffuse light intensity temporal autocorrelation function (g_2_) is recorded in real measurements. From g_2_(r,τ), the normalized diffuse electric field temporal autocorrelation function, g_1_(r,τ), can be extracted by using the Siegert relation, g_2_(r,τ) = 1 + β|g_1_(r,τ)|^2^. Here, β is a constant, y-intercept on the g_2_ axis, depends on the number of spatial modes detected. If the coherence length is sufficiently longer than the photon pathlength, photon detection with a single-mode fiber yields β=0.5 for the unpolarized light. Since g_1_ is the normalized version of the electric field autocorrelation function (G_1_(r,τ)), which has been shown to satisfy the diffuse photon correlation equation based on the partial differential equation [22,23], one can solve this equation analytically or numerically to find G_1 model_(r,τ) and g_1_model_. The dynamic information of the scatterers, here the moving blood cells in the microvasculature, can be obtained by fitting the model to the experimental data. It has been shown that the diffusive motion, αD_B_, can model the dynamics in deep tissue to obtain the blood flow index (BFI), where α is proportional to tissue blood volume fraction, and DB is the effective Brownian diffusion coefficient [21]. The g_1_ function is traditionally fitted to an analytical or numerical solution of the diffusion equation to estimate the BFI parameter. The experimentally relevant parameter, β, is also fitted simultaneously. This approach results in a two-parameter fit performed iteratively by using a nonlinear fitting algorithm (e.g. lsqnonlin in Matlab) which increases the total computational time. This creates a bottleneck, especially when high-speed DCS is used [24] and longitudinal dynamical changes are being investigated at a high data acquisition rate. Therefore, there is a need for fast methods to quantify the BFI parameter. For this aim, we present a fast BFI quantification implementation by using deep learning, as detailed in the next section.

We previously described our DCS system [25], which consists of a continuous-wave laser source (785nm, CrystaLaser, Reno, NV) with a coherence length longer than 10 meters, NIR-optimized single-photon counting modules (SPCM-NIR, Excelitas, Quebec, Canada), and a custom-built eight-channel auto-correlator board (Correlator.com). A multi-mode fiber (1000μm core diameter, FT1000UMT, Thorlabs Inc, Newton, NJ) is used to guide the laser light to the forearm, while single-mode fibers (5μm core diameter, 780HP, Thorlabs Inc, Newton, NJ) are used to collect the emitted light from the forearm to the SPCM at a known distance (Fig. 1). To create blood flow perturbation, a blood pressure cuff was placed on the subject’s forearm. Blood flow data were acquired at 3 stages. First, a baseline measurement is obtained, next, the cuff is inflated to 200 mmHg, finally, it is released. Each stage lasts for approximately 45s.

**FIGURE 1.**
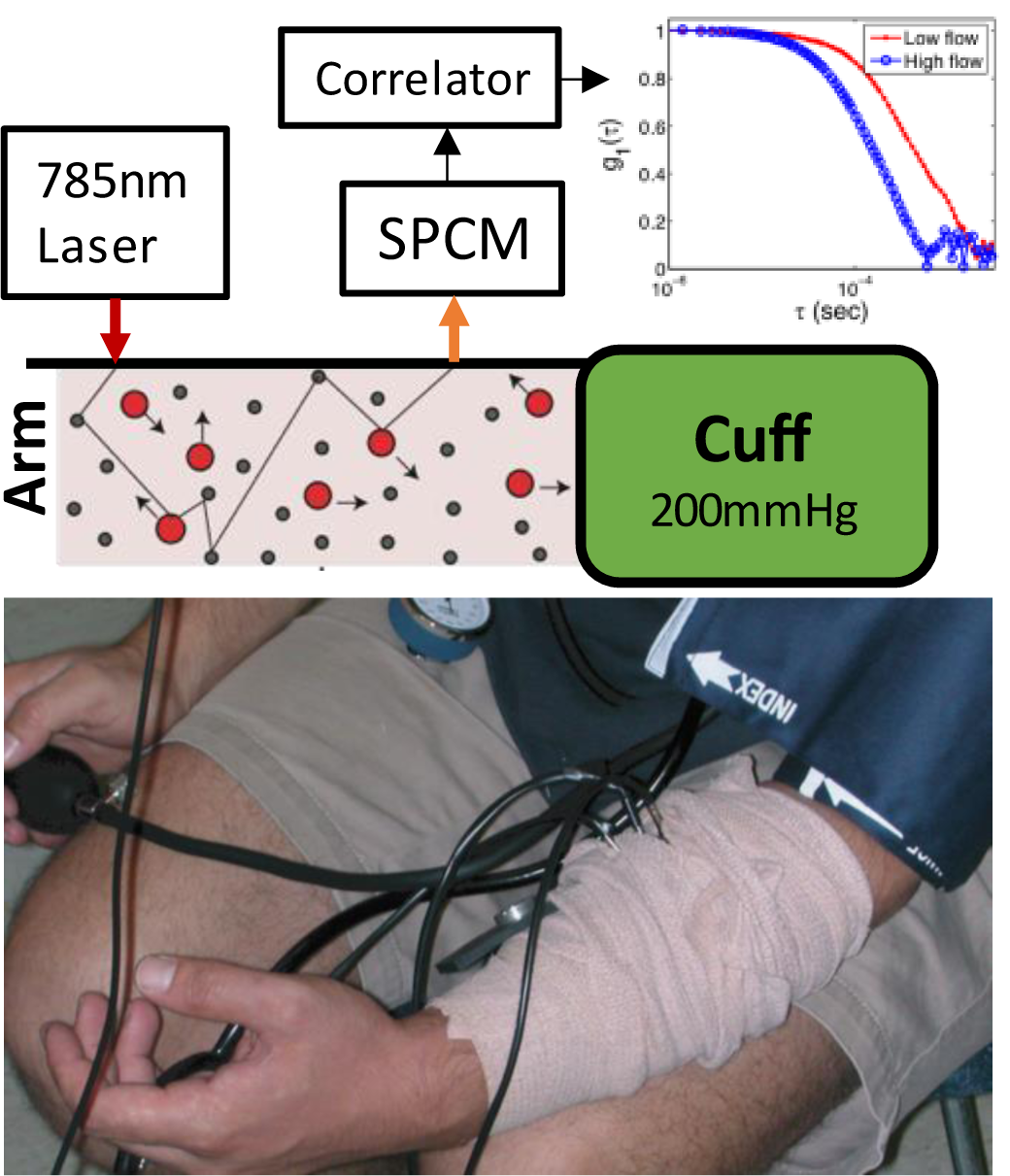
The NIR laser is directed onto the forearm via a fiber. Photons injected by laser light scatter from blood cells, which introduce temporal intensity fluctuations at the single-photon counting module (SPCM) detector. The SPCM detects the photon and sends a signal to the autocorrelator, which generates the g_2_ curve. From g_2_ data, one can derive the g_1_ curve via Siegert relation (see text for details). The decay rate of g_1_ curve is related to blood flow: moving blood cells introduce endogenous flow contrast (sharper decay, higher blood flow). A cuff is also placed on the forearm and pumped to 200mmHg to restrict and perturb the blood flow in the arm muscle microvasculature (low blood flow, slower decay rate (colored red) in g_1_.

### THE DEEP LEARNING NETWORK

We implemented a deep neural network (DNN) to quantify relative blood flow changes, i.e., changes in the blood flow index (BFI, cm^2^/s) as a fast approach. The result of the DNN was also compared to the traditional nonlinear fitting approach. We used the structure of MobileNetV2, which is a lightweight convolutional neural network used for the classification of images [26,27]. We replaced the layers on the bottom and top to accept the measured g_2_ as input and β and BFI as the output parameters. The power of deep learning lies partially in its ability to detect automatically and approximate nonlinear patterns in high-dimensional space [28]. The structure of the DNN is shown in (Fig 2), which uses MobileNetV2 for a total of 161 layers. The model receives an input of 128 g_2_ values that are obtained from the autocorrelator at a specific delay time (τ). It is reshaped into a matrix of 32×4 and fed into a 2D CNN (kernel(1,4), stride(1,4)) with 32 filters, and transposed to map the values into a 32×32 image. The 32×32 image is fed into MobileNetV2 and connected to a fully connected layer of two neurons with a linear activation function that outputs both β and BFI parameters.

**FIGURE 2.**
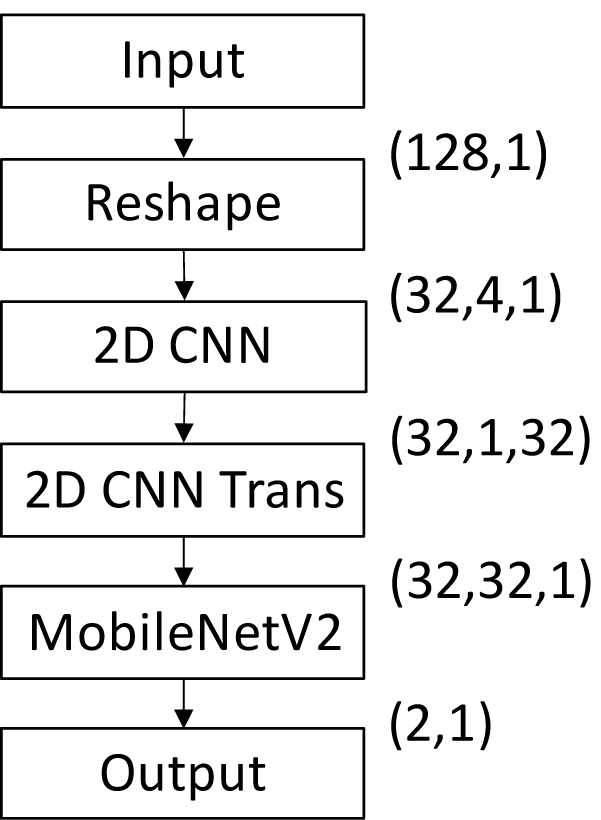
The structure of the deep learning network (DNN). It takes in 128 points of the g_2_ function at a given τ and is reshaped into a 32×4 matrix. Convolutional Neural Network (CNN) is used to map the matrix into a 32×32 image before being passed into MobileNetV2. The output of the MobileNetV2 is connected to a fully connected network of two neurons with a linear activation that gives β and BFI as an output.

The training data were generated by a semi-infinite diffusion model [4], with β sampled from 0.01 to 1 with 120 linearly spaced increments, BFI sampled from 10^−8^ to 10^−5^ cm^2^/s with 120 linearly spaced increments, and DCS noise rate from 10 to 100kcps with 5 linearly spaced increments for a total of 72000 samples. The samples were split into 70% training, 15% validation, and 15% test sets. The DCS noise was obtained as detailed previously [29] and was added to the data to simulate experimental conditions (Fig 3A, C). The multi-tau scheme of a typical auto-correlator, which increases the number of samples averaged as τ increases (hence the smooth tail), has been considered when adding the noise (Fig 3B). The training of the model was performed using Tensorflow 2.0.0 and the mean squared error (MSE) was minimized as a loss function Adadelta was used as the optimizer to adaptively change the learning rate. The training of the model took ∼30.5hr to complete 10000 epochs on a graphics processing unit (Nvidia GeForce GTX1080TI). The details of the training progress after 10000 epochs are shown in Figure 4. The model had a validation mean square error (MSE) of 0.00263 and a training MSE of 0.00067 (Fig. 4A). To check the error against the test set, the predicted value was compared against the target value and the percent error was obtained (Fig 4B). The results show that the predicted β and BFI have 92.3% and 95.6% of all the values within 5% error, respectively. The regression of the trained DNN compared to the test data had an R-square value close to 1, indicating an excellent fit (Fig 4C).

**FIGURE 3.**
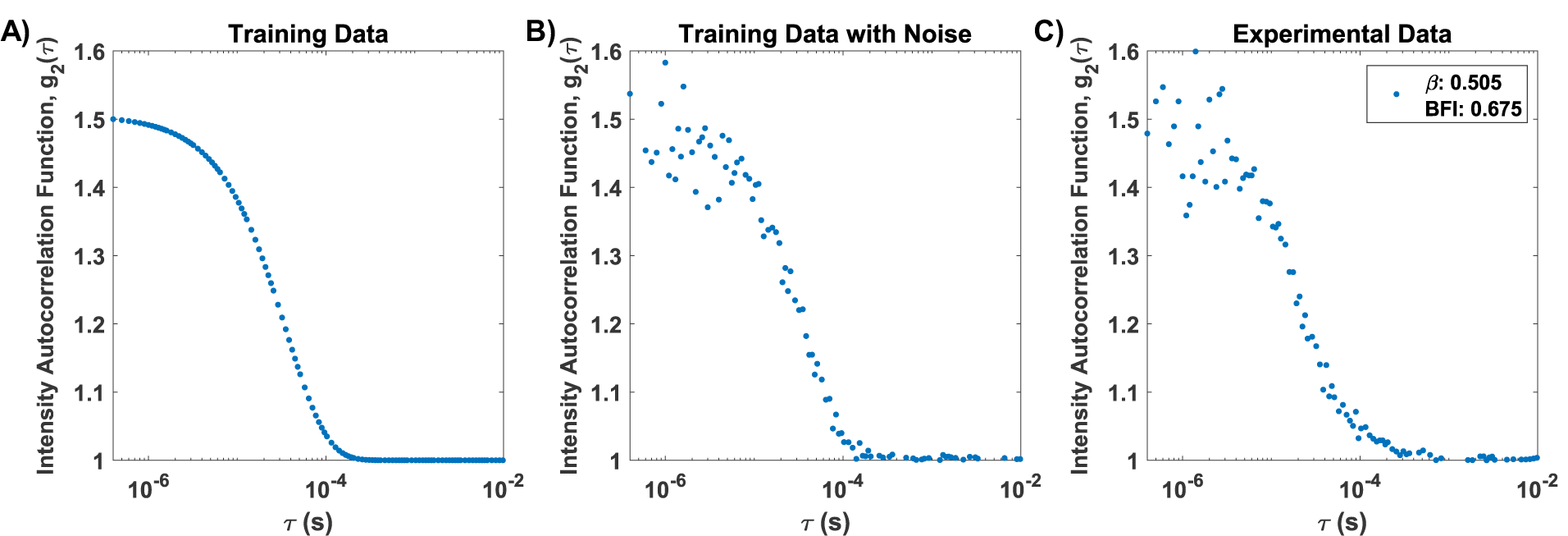
(A) The training curve is generated from the analytical solution. This analytical model is typically used to fit experimental data such as in (C). (B) DCS noise between 10 to 100kcps is added to the training data to simulate experimental conditions. The data is fed into the deep network for training so that it can obtain the correct result even under noisy conditions. (C) This is an example of experimental data.

**FIGURE 4.**
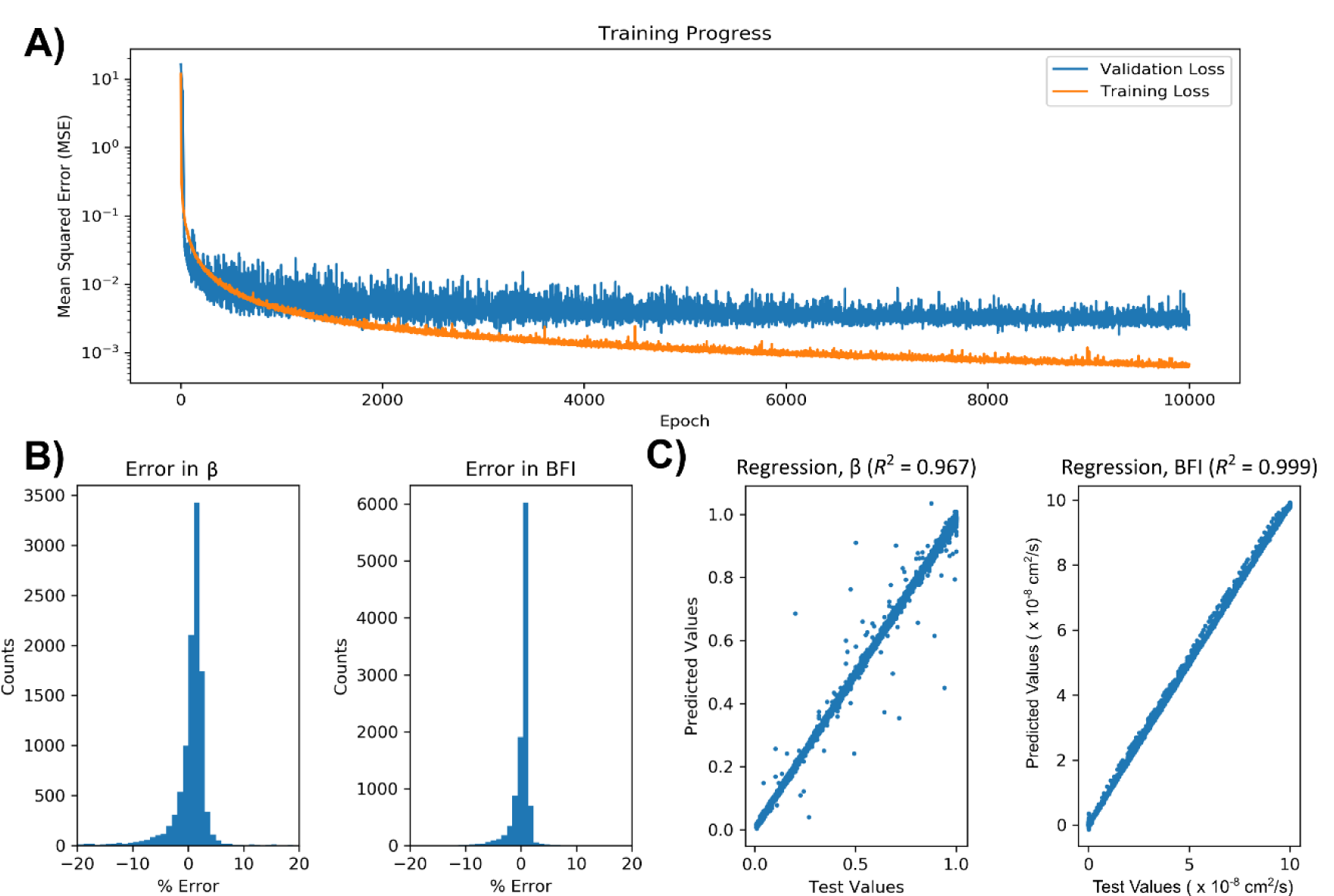
(A) Loss function Mean Squared Error (MSE) after 10000 epochs. The final training MSE and validation MSE is at 0.00067 and 0.00263 respectively. (B) The percent error of the predicted test set as compared to the target value in β and BFI. Here, 92.3% of predicted β and 95.6% of the predicted BFI fall within 5% error from the target. (C) Linear regression of the predicted test set as compared to the target values for both β and BFI both showed good linearity with R2 close to 1.

## RESULT VALIDATION

To validate the model in humans, a pressure cuff experiment was performed to induce occlusion of blood flow in a subject’s forearm (Fig 1). The DCS probe was placed on the subject’s forearm at a source-detector separation of 2.75 cm with an integration time at 1 s. The data were acquired at baseline for 45 s, then the cuff is inflated to induce ischemia for another 45 s, and finally, it was released and allowed to return to the resting state for 45 s. The BFI and β were extracted from both the analytical and the DNN models. As Figure 5 indicates clearly, the DNN showed a good correlation with the analytical solution (AS) for both β and BFI parameters. Both models estimated the β values (Fig. 5A) close to the expected value of 0.5 (AS: 0.488±0.046, DNN: 0.484±0.047) and had the same trend in BFI (Fig. 5B). The root mean square error (RMSE) was calculated to be 0.0573 for β (Fig. 5C) and 0.129 for BFI (Fig. 5D). We also measured the computation time taken to generate each of the 135 data points. The AS fitting method takes 11.19±6.97 ms on each time point to extract BFI and β, while the DNN takes 0.46±0.04 ms on a GPU. This represents a speed increase of more than 2300%.

**FIGURE 5.**
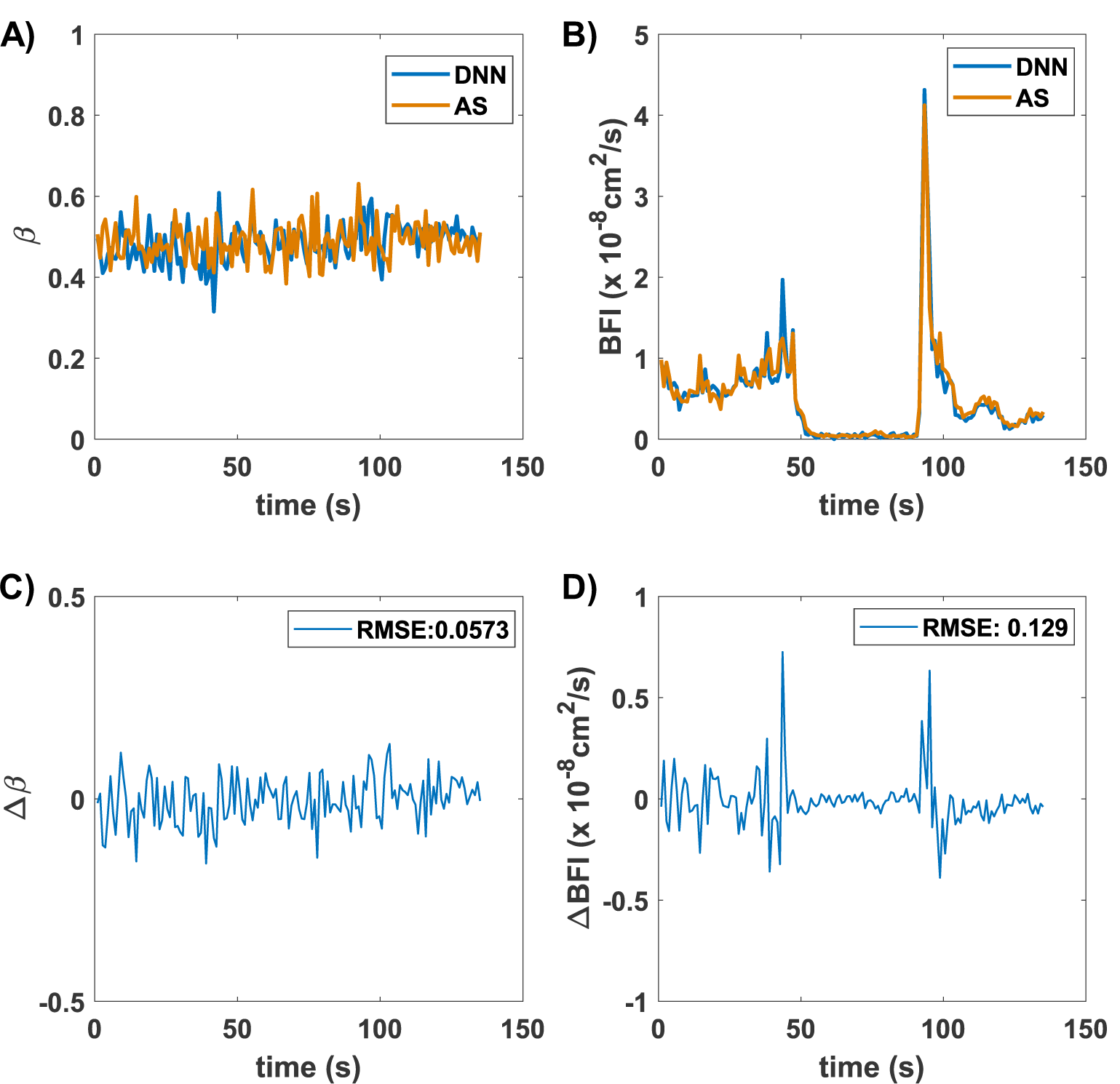
A cuff occlusion experiment was performed by placing the cuff on the forearm and pumped to 200 mmHg to restrict the blood flow. The experiment is split into three blocks of 45 s of rest (baseline), occlusion, and rest again. The analytical solution (AS) was compared to the deep learning model (DNN), which both showed a similar trend for quantified β (A) and BFI (B). The difference between the quantification using AS and DNN was also plotted for β (C) and BFI (D).

## DISCUSSION

For the current feasibility experiment, the integration time was 1 s with a hardware correlator. However, real-time (>=20Hz) software correlators have been introduced recently for fast blood flow monitoring [24], where our approach can show real benefit. For example, for a 20 Hz acquisition rate for a longitudinal monitoring of 10 hours at a neurointensive care unit, the data fit would take 134.4 min vs deep learning approach would take 5.5 min. For the online processing, this represents more than 2 hours of time used for data fitting as compared to a few minutes with DL. Specifically, a 20Hz (50ms integration time), DL can quantify BFI in 0.46 ms while conventional iterative fitting requires 11 ms, which is very significant amount compared to data acquisition time.

For our choice of model, we decided to use a highly optimized DNN that is able to generalize well in different applications. The DNN of choice also has low computing requirements and has a high prediction speed, which is suited for our needs. Previously, we did attempt using a simple multi-layer perceptron (MLP) with limited success. The MLP did present a reasonable level of accuracy (R2 ∼0.95) during regression, but to bring accuracy to a higher level, we have chosen a more complicated yet efficient DNN (MobileNetV2). Additionally, this DNN was also used so that future expansion to quantify more parameters (i.e. quantifying optical parameters) is possible. To show the feasibility and flexibility of DNN, we have trained the DNN with wide ranges in BFI and β values. The choice of the wide range is to show the generalization of the DNN across different scenarios (such as solid or liquid phantom, external light leakages in clinical settings). For deployment in a specific clinical application, the DNN could be trained with a narrower range of BFI and β. This would lead to faster convergence during training and better prediction accuracy of the DNN. The accuracy of the model can be improved by further training the model with experimental data on a phantom with known BFIs. Moreover, the optical properties could also be included in the model by adding two additional neurons to the last layer, which could potentially allow for the estimation of the absolute blood flow and optical parameters within the tissue volume. Alternatively, it is also possible to apply DNN on the raw time-varying signal of photon counts, which may allow for more information to be extracted. This approach allows for the replacement of the autocorrelator with a low-cost acquisition device such as field programmable gate array (FPGA) or a counter/timer. Towards real-time applications, the prediction speed could also be further reduced by pruning weights that are insignificant to reduce the number of calculations needed.

## SUMMARY

Traditionally, a nonlinear curve fitting scheme is used to analyze the DCS data to quantify β and BFI. However, iterative fitting of many data points obtained from dynamic measurements leads to high computation time and cannot provide real-time “online” processing. Thus, a postprocessing step is implemented in the field. In this work, we have shown that deep learning can quantify the BFI and β values from the DCS’s data with high accuracy. This deep learning approach can also increase the quantification speed by more than 2300%. It holds potential towards wide-spread use for clinical applications in the DCS field, especially for those that require longitudinal, fast, real-time monitoring of dynamic changes, such as monitoring of muscle and brain function studies.

## ACKNOWLEDGMENT

The authors would like to acknowledge financial support from the Ohio Third Frontier to the Ohio Imaging Research and Innovation Network (OIRAIN, 667750).

The authors declare no conflicts of interest.

